# Irrelevant Predictions: Distractor Rhythmicity Modulates Neural Encoding in Auditory Cortex

**DOI:** 10.1101/2020.01.27.920728

**Authors:** Shiri Makov, Elana Zion-Golumbic

## Abstract

Dynamic Attending Theory suggests that predicting the timing of upcoming sounds can assist in focusing attention towards them. However, whether similar predictive processes are also applied to background noises and assist in guiding attention *away* from potential distractors, remains an open question. Here we address this question by manipulating the temporal predictability of distractor sounds in a dichotic listening selective attention task. We tested the influence of distractors’ temporal predictability on performance and on the neural encoding of sounds, by comparing the effects of Rhythmic vs. Non-rhythmic distractors. Using Magnetoencephalography (MEG) we found that, indeed, the neural responses to both attended and distractor sounds were affected by distractors’ rhythmicity. Baseline activity preceding the onset of Rhythmic distractor sounds was enhanced relative to Non-rhythmic distractor sounds, and sensory response were suppressed. Moreover, when distractors were Rhythmic, responses to attended sounds were more strongly lateralized to the contra-lateral hemisphere. Behavioral performance also improved in the Rhythmic condition. These combined behavioral and neural results suggest that not only are temporal predictions formed for task-irrelevant sounds, but that these predictions bear functional significance for promoting selective attention and reducing distractibility.

---

Auditory attention is the act of selecting task-relevant sounds and processing them preferentially relative to background stimuli. One way that attentional selection manifests in the auditory system is through enhanced neural responses to attended sounds as compared to unattended sounds (Hillyard et al. 1973; Woldorff et al. 1993; Ross et al. 2004; Atiani et al. 2014; Fuglsang et al. 2017; Jaeger et al. 2018; Fiedler et al. 2019). A prerequisite for this selective enhancement is knowing or predicting the sensory attributes of to-be-attended sounds, such as pitch, timing and location. These predictions can be used to modulate neural responses within the specific neuronal populations coding these attributes, through gain-changes or receptive-field sharpening (Fritz et al. 2007; Jaramillo and Zador 2011; Lakatos et al. 2013; Lu et al. 2017), as suggested by the predictive coding framework (Sherwell et al. 2017; Heilbron 2018; Gordon et al. 2019). There is also evidence suggesting that having explicit knowledge regarding the sensory attributes of *task-irrelevant* sounds (distractors), can reduce their negative consequences on performance and modulate neural responses (Melara et al. 2002a; Chait et al. 2010; Couperus and Mangun 2010; Melara et al. 2012; Noonan et al. 2016). This suggests that similar principles for guiding attention through prediction may also operate on stimuli outside the ‘spotlight’ of attention (Noonan et al. 2018). However, how the predictability of distractor sounds ultimately impacts selective attention and interacts with neural encoding of both attended and distractor sounds themselves, remains highly unknown.

One domain where predictability can be particularly beneficial for selective attention is timing (Nobre and Coull 2010; Zion Golumbic et al. 2012a; Lakatos et al. 2013; Nobre and Ede 2017). Being able to anticipate *when* sounds will occur is extremely instrumental for actively guiding perception and attention, as put forth by Dynamic Attending Theory (Large and Jones 1999; Cason and Schön 2012; Brochard et al. 2013; Cravo et al. 2013; Jones 2019). Predictions stemming from temporal regularities within continuous sound sequences – such as isochronous or syncopated rhythms – are of particular interest since they need to be inferred from the stimulus itself, without explicit instructions (Escoffier et al. 2015; Sohoglu and Chait 2016; Tal et al. 2017; ten Oever et al. 2017; Wollman and Morillon 2018). Rhythms are also prevalent in many natural stimuli such as speech, music and biological motion, making the ability to form and utilize temporal predictions ecologically relevant for real-life perception and attention (Haegens and Zion Golumbic 2018; Rimmele et al. 2018). Mechanistically, it has been proposed that selective attention to rhythmic stimuli harnesses the inherent oscillatory properties of neuronal populations for adjusting population excitability in a temporally-precise manner, making for a metabolically efficient implementation of temporal predictions (Schroeder and Lakatos 2009; Nobre and Ede 2017).

The substantial evidence for the utility of the temporal predictability for facilitating selective attention to rhythmic sequences of sounds begs the question whether temporal predictions are also derived for rhythmic stimuli that are task-irrelevant. And if so, are these predictions utilized to promote behavioral goals? Indeed, the auditory system does seem to be sensitive to the timing of unattended sounds, as reflected, for example, by Mismatch Negativity (MMN) responses to interval and duration deviants (Lyytinen et al. 1992; Escera et al. 2002; Winkler et al. 2003; Kisley et al. 2004; Zion-Golumbic et al. 2007; Lumaca et al. 2018; Thomassen and Bendixen 2018). Besides being important for auditory stream segregation (Nelken 2004; Näätänen et al. 2010; Thomassen and Bendixen 2018), forming representations for the temporal structure of distractor sounds also seem to bear a functional role in facilitating selective attention (Sussman 2017). For example, several behavioral studies have shown that performance on selective-attention auditory tasks was less disrupted by the presence of rhythmic vs. random auditory distractors (Devergie et al. 2010), at least in young adults (Rimmele et al. 2012) and when the task was sufficiently difficult (Andreou et al. 2011). Moreover, Escoffier et al. (2010) showed that reaction times on a visual discrimination task were facilitated at moments corresponding to the beat of a task-irrelevant background rhythm. These findings suggest that not only is the timing of irrelevant rhythmic auditory sequences encoded, but that it can be beneficial for withstanding distraction and improving performance.

However, how this facilitatory effect is implemented in auditory cortex, is unknown. Specifically, are cortical responses to distractor sounds enhanced or suppressed when they are temporally predictable? And in what way does the temporal structure of distractor sounds influence neural responses to attended sounds? The current study attempts to address these questions, by measuring neural responses to attended and distractor sounds in a dichotic listening selective-attention task. We recorded brain activity from human participants using Magnetoencephalography (MEG) and tested the effects of Rhythmic vs. Non-rhythmic auditory distractors. We equated the global temporal statistics in the two distraction conditions, and in order to avoid confounds due to low-level sensory differences, we also carefully controlled the local acoustic coincidence between the attended and distractor sound sequences. This enabled us to explore how the temporal predictability of distractors affects the way attended and distractor sounds are represented in cortical activity, as well as the effect it exerts on behavioral performance.

## Materials and Methods

### Participants

21 participants (12 female, mean age 25.0±4.6, range 19-37), with self-reported normal hearing and no history of neuropsychiatric disorders, took part in this study. One participant was excluded from the MEG analysis for lack of sufficient data. Two additional participants were excluded from the behavioral analysis due to technical reasons. The study was approved by the Institutional Ethics Committee at Bar Ilan University. Participants provided their written consent prior to commencement of the experiment and were paid for their participation.

### Stimuli and Task

Two auditory streams were delivered dichotically via MEG compatible in-ear tubephones (ER3-A, Etymotic Research), adjusted to a comfortable listening level. Subjects were instructed to attend to the sounds presented to the left ear, and to ignore distractor sounds presented to the right ear (Fig. 1A). This ear-pairing remained constant across all conditions and participants in order to maximize the number of trials in each condition and to allow lateralization analysis. The to-be-attended stream consisted of an oddball sequence of pure tones presented at a constant rate of 1.6 Hz. Standard tones were 500 Hz (30 ms, including 5 ms raise and fall time), and target tones (∼13%) were slightly lower in pitch (430-480 Hz, set individually to be just above threshold; see below). Participants were instructed to respond to target tones via button press.

**Figure 1:**
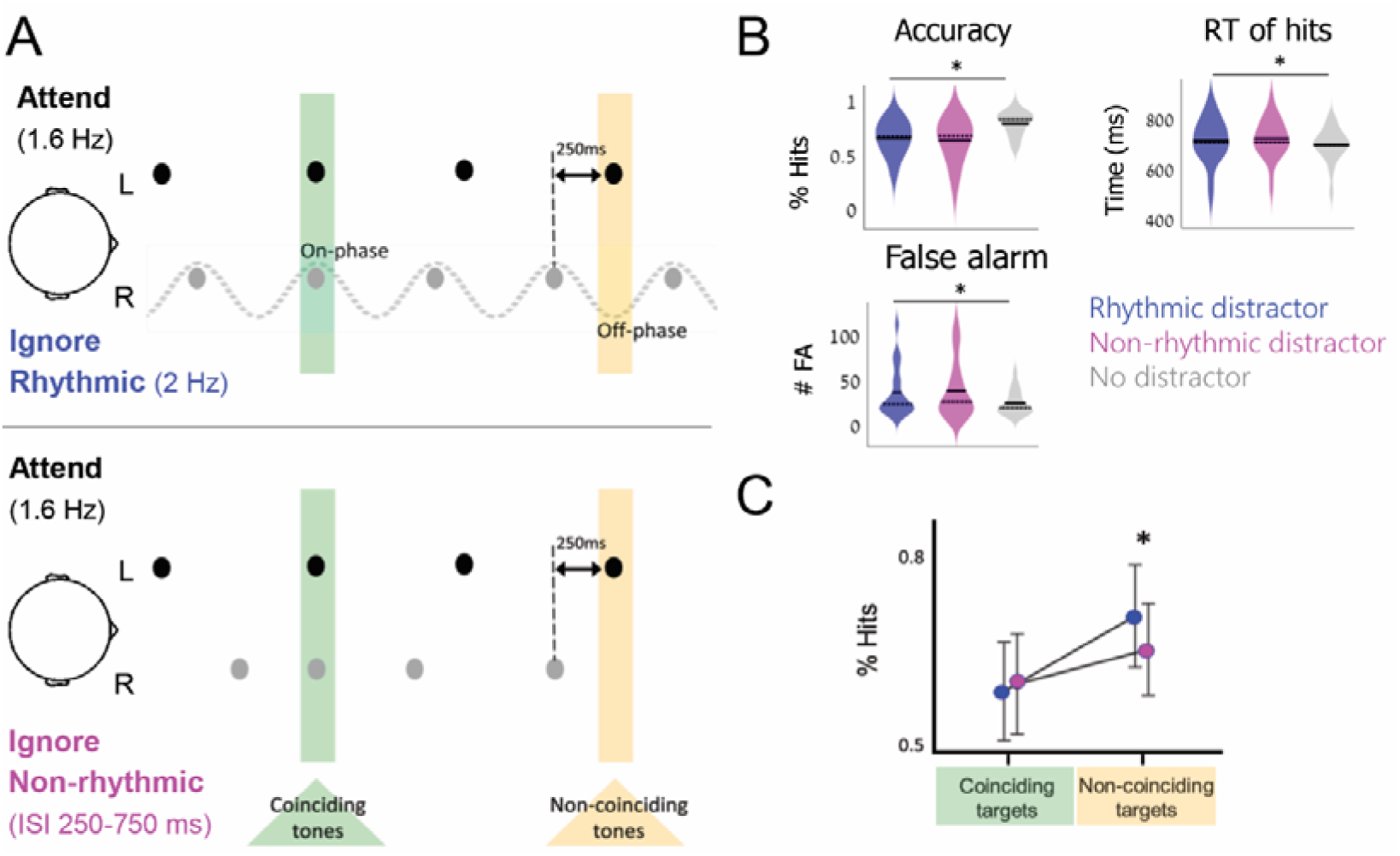
Experimental design and behavioral results. (A) The temporal relation between sounds in the to-be-attended (left ear, black dots) and to-be-ignored (right ear, gray dots) streams. The to-be-attended stream consisted of an oddball sequence of tones presented at a fixed rate of 1.6 Hz (SOA 625 ms; 500 Hz standards, slightly above threshold pitch deviants). The to-be-ignored (distractor) stream consisted of a sequence of 680 Hz pure tones that was either Rhythmic (top; 2 Hz, SOA 500 ms) or Non-rhythmic (bottom; SOA 250-750 ms). The temporal relations between the attended (black) and ignored (gray) tones were controlled such that in both distraction conditions 25% of the attended tones coincided with a distractor tone (“coinciding tones”, green) and 25% appeared 250 ms following a distractor (“non-coinciding attended tones”, yellow). Target sounds occurred only at these positions, ensuring control of low-level acoustics between the Rhythmic and Non-rhythmic Distraction conditions. (B) Accuracy, False alarm rates and RT in response to targets appearing in the presence of Rhythmic (blue), Non-rhythmic (pink) or No distraction (gray). In each violin plot (Hoffmann 2015), the full and dashed horizontal lines denote the mean and median, respectively. All behavioral measures indicate that both distraction types were disruptive for task performance. (C) Modulation of target-detection accuracy by Distraction Type (Rhythmic vs. Non-rhythmic) and Temporal Coincidence between the streams (Coinciding vs. Non-coinciding). Accuracy was improved for targets occurring 250 ms after a distractor tone in the Rhythmic condition (namely, at the off-phase of the distractor rhythm).

The to-be-ignored stream consisted of pure tones (680 Hz), presented either Rhythmically at a rate of 2 Hz, or in a Non-rhythmic fashion. Importantly, SOAs in the Non-rhythmic distractor stream were carefully constructed to match the temporal statistics of the Rhythmic distractor, both at a global and local scale, in the following ways: (1) Rhythmic and Non-rhythmic distractor streams consisted of exactly 4 tones in every 2-second interval, yielding identical mean-rates of 2 Hz; (2) 25% of the attended tones occurred at the same time as distractor tones (“Coinciding tones”; Fig. 1A, green) and 25% of attended tones occurred exactly 250 ms after distractor tones (“Non-coinciding attended tones”; Fig. 1A, yellow). This ensured comparable local acoustic in the Rhythmic and Non-rhythmic Distraction conditions around both coinciding and non-coinciding attended tones. Moreover, target tones were strategically placed only at these coinciding or non-coinciding positions relative to the distractor stream, making the behavioral responses comparable across conditions. (3) Similarly, we equated the number of distractors that occurred exactly 250 ms after an attended sound in the Rhythmic and Non-rhythmic Distraction conditions (“non-coinciding distractor tones”; Fig. 4A). This further allowed us to avoid low-level confounds when assessing the effects on neural responses to distractor sounds. Accordingly, all event-related analyses were restricted to these subsets of non-coinciding sounds, where the local acoustics were perfectly matched across the Rhythmic and Non-rhythmic Distraction conditions.

The experiment consisted of 30 trials, lasting 65 seconds each. Each trial began with the presentation of 12 tones to the to-be-attended ear (7.5 seconds, including 2 targets), to assist participants in focusing attention before the distractor stream started. This initial epoch was excluded from all analyses, as well as additional 5.1 seconds after the distractor streams started to appear, to allow sufficient time for establishing the temporal structure of the distractor. Thus, the analysis focused on the latter 52.4 seconds of each trial. Besides the Rhythmic and Non-rhythmic Distraction conditions, an additional No-distraction control condition was included, where only attended sounds were presented. In all trials, participants were instructed to fixate the center of a gray screen where the word “listen” appeared. The total duration of the task was 38.2±2.1 (range 35.1-42.7) minutes.

### Personalized threshold procedure

To ensure that the target-detection task was sufficiently challenging, the pitch of target tones was set individually to be just above detection threshold related to standard tones in the attended stream. For that aim, participants underwent a staircase procedure prior to the main experiment, to determine their personal pitch-difference threshold (10.3±4.9, range 5-22 minutes). The staircase procedure involved a similar oddball task as used in the main experiment in the No-distraction condition, and was performed while lying in the MEG. Thus, participants performed a deviance-detection task on a tone-sequence presented to the left ear, and the pitch of the target was adapted on-line using a 2-down 1-up procedure (Levitt 1971) with step-size of 1 Hz. The staircase procedure terminated after three consecutive direction shifts (up-down-up) and subsequent testing verified target detection rates between 76-86% (when no distractor was present).

### MEG Data acquisition

MEG data was acquired in supine position, using a whole-head 248-channel magnetometer array (4-D Neuroimaging, Magnes 3600 WH) in a dimly and magnetically shielded room. Data were sampled at 1017.23 Hz and bandpass-filtered online at 0.1 to 400 Hz. Head position in relation to the MEG sensors was estimated using 5 coils that were attached to the participant’s scalp. Head shape was digitized manually for later source estimation.

### MEG data preprocessing and source estimation

MEG data preprocessing was performed in MATLAB (The MathWorks) using the FieldTrip toolbox (Oostenveld et al. 2011) and custom written scripts. External noises (power-line, mechanical vibrations of the building), as well as heart-beat artifacts, were removed offline using a predesigned algorithm (Tal and Abeles 2013). We excluded from all data the first 12.6 seconds of each trial (see *Stimuli and Task*), to avoid introducing onset effects and allow sufficient time for establishing the temporal structure of the distractor. Independent component analysis (ICA) was used for removal of blinks, eye movements and residual heartbeat traces. Remaining outlier trials (4.2±4.4%, range 0.4-13.2%) were identified manually by visual inspection and were discarded from event-related analysis, however all trials were included in the evoked power analysis (see below).

In order to assess the neural source of MEG-recorded activity, we applied source estimation analysis on the preprocessed data, using the MNE-python platform (Gramfort 2013; Gramfort et al. 2014), and additional custom written Python scripts (www.python.org).

Source modeling was performed by computing Minimum-Norm Estimates (MNEs; Hämäläinen and Ilmoniemi 1994). For calculating the forward solution, we constructed a Boundary Element Model (BEM) for each subject based on head shape and location relative to the MEG sensors, and a structural MRI template (average brain of FreeSurfer software; http://surfer.nmr.mgh.harvard.edu/). BEM was used to constrain MEG source locations to the cortical surface. The cortical surface of each participant was decimated to 8196 source locations per hemisphere with at least 5-mm spacing between adjacent locations. A noise covariance matrix was estimated using sections of the data where no stimuli were presented, taken from a 250 ms pre-stimulus period in the No-distraction condition (averaged over 106.3 ± 37.1 epochs, range 62-211). Activity at each source location was estimated using an inverse operator, computed from the forward solution and the noise covariance matrix. For visualizing the current estimates on the cortical surface, we used dynamic Statistical Parametric Map (dSPM), which is an F-statistic indicating locations where the MNE amplitudes are above the noise level (Dale et al. 2000). To compensate for inter-subject differences, individual cortical surfaces were morphed to a common space through spherical surface into FreeSurfer average brain, with 10,242 dipoles per hemisphere (Fischl et al. 1999). The dSPM time courses and power spectrum were extracted from 20 predefined cortical regions of interest (ROIs; 10 per each hemisphere) across the lateral cortical surface in each hemisphere. These included: Early Auditory Cortex, Auditory Association Cortex, Posterior Opercular Cortex, Anterior-Inferior Parietal Cortex, Anterior-Superior Parietal Cortex, Temporal-Parietal Junction, Insular and Frontal Opercular Cortex, Orbital and Polar Frontal Cortex, Dorsolateral Prefrontal Cortex, Inferior Frontal Cortex. These ROIs were identified anatomically according to the Human Connectome Project multi-modal parcellation atlas (Glasser et al. 2016) (see Figure 4).

### Data Analysis – Behavior

Responses to target-tones occurring within a window of 150-1250 ms after target onset were considered “hits”. All other responses were considered false alarms (FAs). Reaction times (RTs) were assessed only for “hit” responses. To test the accuracy and RT measures, we fitted generalized linear mixed effects models, implemented in the statistics software R (Team and R Development Core Team 2016) using the lme4 (Bates et al. 2015) and lmerTest (Kuznetsova et al. 2017) packages. For fitting accuracy rates we used a logit link function, with p-values based on asymptotic Wald tests that are provided in the summary of R’s glmer function. For the RT model, p-values were obtained using the Satterthwaite’s degrees of freedom approximation (Satterthwaite 1946). All models converged with a maximal random structure, as random intercepts and slopes of all predictors were included, adjusted by subjects.

For each behavioral measure (accuracy and RT) we fitted a model to test the overall performance, using the Temporal Structure of the distractor as predictor. This predictor was Helmert coded: one beta codes for the contrast between the No-distraction condition and the average in the two Distraction conditions, and the second beta codes for the contrast between The Rhythmic and Non-Rhythmic distraction conditions.

We then applied a second model on the hit-rates data, to explore the interaction between the distractors’ Temporal Structure (Rhythmic vs. Non-rhythmic) and the Temporal Coincidence of targets and distractor tones (Coinciding vs. Non-coinciding), using both variables and their interaction as predictors. Variables were sum coded, i.e. each beta reflects the difference from the grand mean. The non-balanced number of “hits” in each category precluded similar analysis of RTs.

The measure of FA rates included only one value per condition for each subject, and thus was statistically assessed via permutation tests, contrasting (a) No-distraction vs. both Distraction conditions and (b) Rhythmic vs. Non-rhythmic distraction.

### Data Analysis - MEG

Two types of analyses were applied to MEG data, looking at: (1) the evoked steady state response (SSR) to the attended stream (1.6Hz) and (2) event-related field (ERF), time-locked to the onset of both attended and distractor tones. These were performed both at the scalp-level, and at the source level. Analyses at the scalp-level were performed using the FieldTrip toolbox and custom-written scripts in Matlab. Analyses of source-level were performed using MNE-python platform (Gramfort 2013; Gramfort et al. 2014), with in-house written scripts in Python (www.python.org). This two-tiered approach, examining the data at both the scalp and source levels, provides complementary perspectives for looking at the recorded neural responses.

#### Evoked Steady-State Response Analysis (SSR)

The evoked-power analysis was aimed at extracting the magnitude of the steady state response (SSR) that was evoked by the attended sequence, presented at a rate of 1.6 Hz. Importantly, this attended stimulus was identical in all conditions, and therefore any observed modulation of the 1.6 Hz SSR should be attributed to the existence and type of the concurrently presented distractor. The SSR was quantified by averaging all 10 full-length trials (52.4 second) within each condition and each participant and applying a Fast Fourier Transform (FFT; Hanning window) to this time-locked data. We tested for two planned contrasts, comparing the SSR: (I) under No-distraction vs. the average across the two dichotic conditions and (II) under Rhythmic vs. Non-rhythmic distraction. For testing these planned contrasts at the scalp level, paired t-tests were performed at each MEG sensor individually (threshold α=0.05), and corrected for multiple comparisons across sensors using a cluster-based permutation procedure to determine whether the observed cluster size falls in the top 5 %^tile^, relative to a null distribution (10,000 permutations, statistic: T-value sum within cluster). The permutation test was performed separately over each hemisphere, based on an a-priori assumption that responses may vary between hemispheres based on their ipsi/contra lateral status relative to the presentation of each sound (Sassenhagen and Draschkow 2019). For source-level analysis, power at 1.6 Hz was averaged in each predefined ROIs (see above). Paired t-tests were applied within each region. At the source level we report both uncorrected p-values within each ROI as well as fdr corrected p-values for multiple comparisons across regions.

#### Event-Related Field Analysis (ERF)

We further tested the MEG response to individual tones using ERF analyses. We time-locked the neural response separately to two types of tones: (I) Attended non-coinciding tones, (II) Distractor non-coinciding tones. Given the careful design of the stimuli, each of these tones were perfectly matched in their low-level acoustic properties in the Rhythmic and Non-rhythmic conditions, and therefore any differences between them can be reliably attributed to the temporal structure of the distractor. Target tones, as well as all tones occurring within 700 ms after them, were excluded from this analysis to prevent residual noise from motor responses. For each type of tone (Attended or Distractor) we equated the number of trials from the Rhythmic and Non-rhythmic Distraction conditions by randomly sub-sampling (number of trials: Attended non-coinciding tones: 98.3±36.9, range 49-201; Distractor non-coinciding tones: 129.6±27.7, range 77-201). Trials were epoched between −100 and 250 ms around the onset of each tone and the ERF was calculated for each subject by averaging the response across trials.

Statistical analysis focused on potential effects of distractor-rhythmicity on the ERFs to both Attended and Distractor tones. This was carried out at the scalp and source levels, using a series of paired t-tests comparing the Rhythmic vs. Non-rhythmic distraction conditions, in non-overlapping 20 ms-long time windows, from −40 to 240 ms after tone onset.

At the scalp-level, t-tests were performed at each MEG sensor individually α =0.05) and corrected for multiple comparisons using a cluster-based permutation test separately for each hemisphere (10,000 permutations, statistic: T-value sum within cluster), followed by Bonferroni correction for multiple comparisons over the time-windows tested. At the source-level, t-tests were performed on the average within each of the 20 predefined ROIs (see above), and corrected for multiple comparisons using spatial-temporal clustering (10,000 permutations, statistic: T-value sum within spatial-temporal cluster) separately per hemisphere.

## Results

### Behavioral results

The difficultly of the target-detection task was tailored-individually so that targets had a just-above-threshold pitch difference, to avoid ceiling effects. Indeed, mean hit rates across all conditions were around 65.3±20.2% (range 28.9-85.1% across participants), indicating that the task was not trivial (Fig. 1B, left).

Comparisons of accuracy and RTs across conditions were done using generalized linear mixed effects models (see *Methods*). These tests revealed that both accuracy and RTs were improved in the No-distraction condition relative to the two dichotic conditions (accuracy: β=0.79±0.18, z=5.97, p<0.001; RT: β=−24.27±6.61, t=−3.67, p=0.002; Fig. 1B), indicating that the presence of distractor sounds of any type interfered with performance. In the same direction, FA responses were more frequent under distraction rather than in the No-distraction condition (p<0.001). No significant differences were found between the two Distraction Types on accuracy (β=-0.07±11, z=-0.66, p=0.51, n.s.), RT (β=8.17±6.32, t=1.29, p=0.21, n.s.) or FA (p=0.8, n.s.), when pooling together all targets.

However, since in the current design targets were strategically placed to either coincide with a distractor tone or to occur exactly 250 ms after a distractor (Fig. 1A), we conducted an additional analysis, testing for the effect of this Temporal Coincidence on performance. We compared the effects of Distractor Types (Rhythmic vs. Non-rhythmic) x Temporal Coincidence (Coinciding vs. Non-coinciding) on accuracy using a mixed effects model. There was a significant interaction between these two factors (β=-0.08±0.03, z=-2.45, p=0.014; Fig. 1C), indicating that detection accuracy of Non-coinciding targets was improved when the distractor was Rhythmic vs. Non-rhythmic, whereas for targets Coinciding with a distractor there was no effect of Distractor Type.

### MEG results

#### Evoked Power: Steady-State Response to Attended Sounds

Task-relevant tones were presented to the left ear at a constant rate of 1.6 Hz in all experimental conditions (Fig. 1A). Isochronous stimuli are known to elicit a steady-state response (SSR) at the rate of their presentation that is clearly visible as a peak in the evoked power spectrum (Regan 1966; and Fig. 2A). We tested whether this SSR was affected by the presence and temporal structure of distractor tones presented concurrently to the other ear. To this end, we performed two planned contrasts. Comparing the SSR under No-distraction vs. the average of the two dichotic conditions at the scalp-level, revealed that the SSR was significantly stronger when there was No-distraction, particularly over the ipsilateral (left) hemisphere (Fig. 2 B-D; cluster permutation test p<0.001). Testing at the source-level showed that this ipsilateral effect is localized at tempo-parietal areas (Posterior Opercular Cortex and anterior-inferior Parietal Cortex, paired t-test p=0.006 and p=0.005, respectively, fdr corrected; Fig. 2D). No significant effects were observed in the contralateral (right) hemisphere (scalp-level p=1, source-level: all p>0.09). Thus, the presence of concurrent distractor tones seemed to have biased the attended response to be more lateralized to the contralateral hemisphere, relative to when there was no distraction.

**Figure 2:**
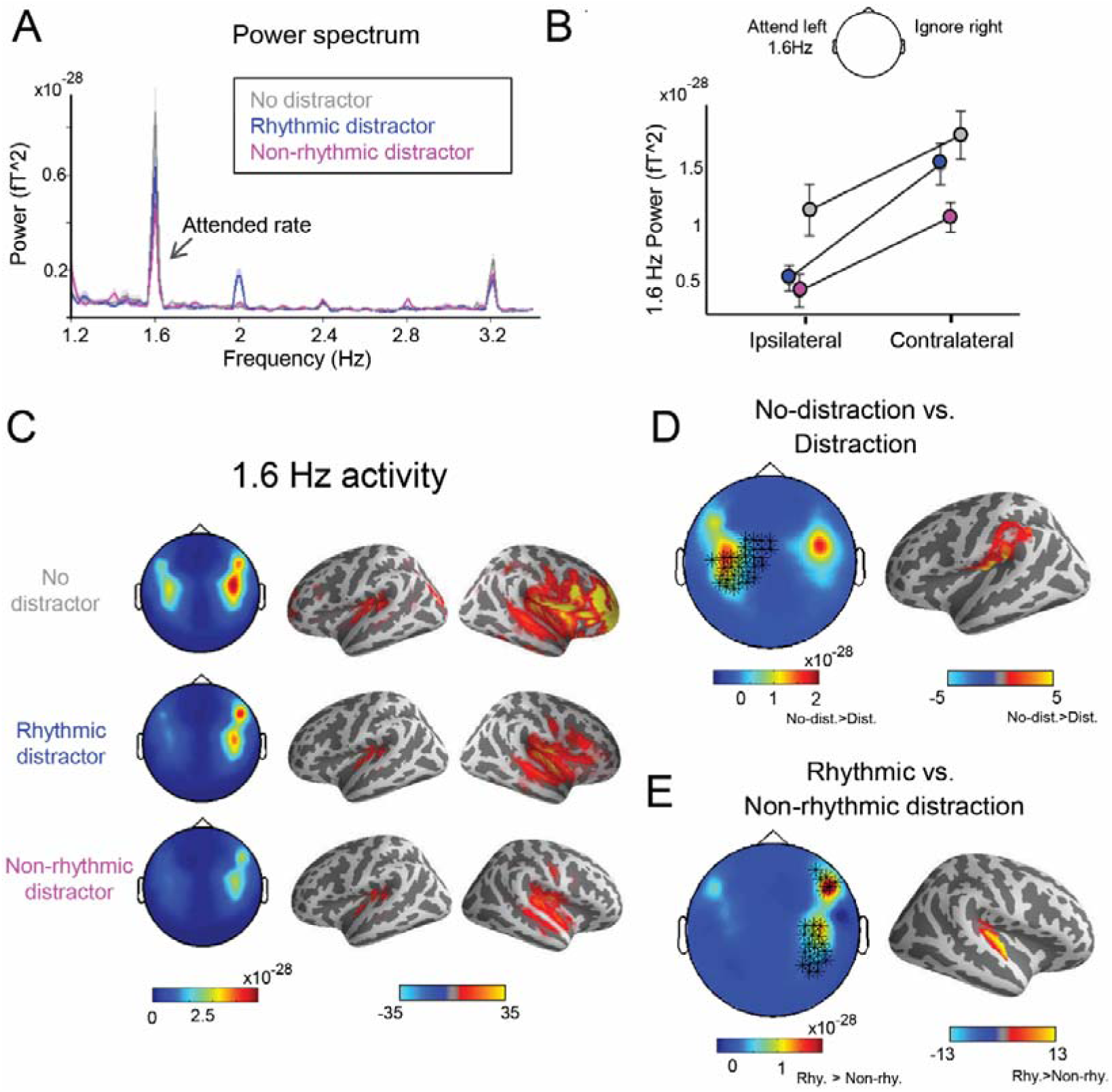
Spectral analysis results. (A) Evoked power spectrum averaged across all MEG sensors demonstrating a peak at the attended rate (1.6 Hz) representing the SSR in all three conditions (gray - No-distraction, blue - Rhythmic distraction and pink - Non-rhythmic distraction). Shaded areas denote standard error of the mean (SEM). (B) The magnitude of the SSR, averaged across all sensors over the ipsilateral (left) and contralateral (right) hemispheres. SSR was consistently stronger in the hemisphere contralateral to the stimuli, and this lateralization was strongest in the presence of a concurrent Rhythmic distractor. (C) Scalp-level topographies (left) and source estimation (right) of the 1.6 Hz SSR in the three experimental conditions. (D) Subtracted topographies depicting the difference in SSR in the No-distraction condition vs. the average across the two dichotic conditions. Left: scalp-level data. Asterisks denote MEG channels in which the contrast yielded a significant difference (α=0.05, cluster corrected). Right: the estimated source the effect found in the ipsilateral hemisphere, localized to left Posterior-Opercular Cortex and the anterior portion of the Inferior-Parietal Cortex. (E) Subtracted topographies depicting the difference in SSR in the Rhythmic vs. Non-rhythmic Distraction conditions. Left: as in D. Right: the estimated source the effect found in the contralateral hemisphere, localized to right Early Auditory Cortex.

The second comparison, contrasting the SSR under Rhythmic vs. Non-rhythmic distraction, revealed that the response in the contralateral (right) hemisphere was stronger in the presence of Rhythmic distractors (Fig. 2E; cluster permutation test p<0.001). Statistical testing at the source-level showed that this enhanced response was localized in the contralateral (right) Early Auditory Cortex, (paired t-test p=0.044, uncorrected; Fig. 2E), with a trend in Association Auditory Cortex (paired t-test p=0.076, uncorrected), although the statistical significance of this effect did not survive correction for multiple comparisons across ROIs. No significant effects were observed in the ipsilateral (left) hemisphere (scalp-level p=1, source-level: all p>0.13). Importantly, spectral analysis of the combined auditory stimulus in the different conditions confirmed that the observed modulations in the SSR at 1.6Hz cannot be trivially attributed to differences in 1.6Hz power in the acoustic input between condition.

We found no significant Pearson correlations between the magnitude of the 1.6 Hz SSR and behavioral outcomes (hit rates, false alarm rates, mean RT or RT variability; all ps>0.07, uncorrected).

#### Event-Related Analysis: Non-Coinciding Attended Tones

Next, we tested how the temporal structure of distractor sounds affected neural responses to individual tones, by looking at time-locked ERF responses. For Attended tones, this analysis focused on responses to standard tones occurring precisely 250 ms after a distractor (Attended Non-coinciding tones, see *Methods* and Fig. 1A). As described above, these epochs were perfectly controlled for local low-level acoustics across the Rhythmic and Non-rhythmic conditions, therefore any differences in the ERFs can be reliably attributed to the temporal structure of the distractors.

The scalp-level ERFs and their projection onto source space are presented in Fig. 3. Scalp-level ERFs to attended tones displayed a 4-pole scalp topography typically observed for MEG-recorded auditory evoked responses, with a characteristic M1 response at ∼100ms. This response was significantly modulated by the temporal structure of the distractors, however only on the left (ipsi-lateral) side (cluster p<0.001, Bonferroni corrected; Fig. 3B). These ipsi-lateral M1 responses to (non-coinciding) attended sounds were reduced when distractors were Rhythmic. Source estimation of the ERFs mapped onto auditory areas. In line with the scalp-level results, a trend was observed in ipsi-lateral (left) auditory cortex, showing reduced M1 when distractors were Rhythmic, although this did not reach statistical significance (p=0.11 uncorrected; Fig. 3C). We found no evidence for such modulation of attended ERF by the type of distractor in the contra-lateral (right) hemisphere (scalp-level: p=0.09; source-level: p=0.53; Fig. 3D).

**Figure 3:**
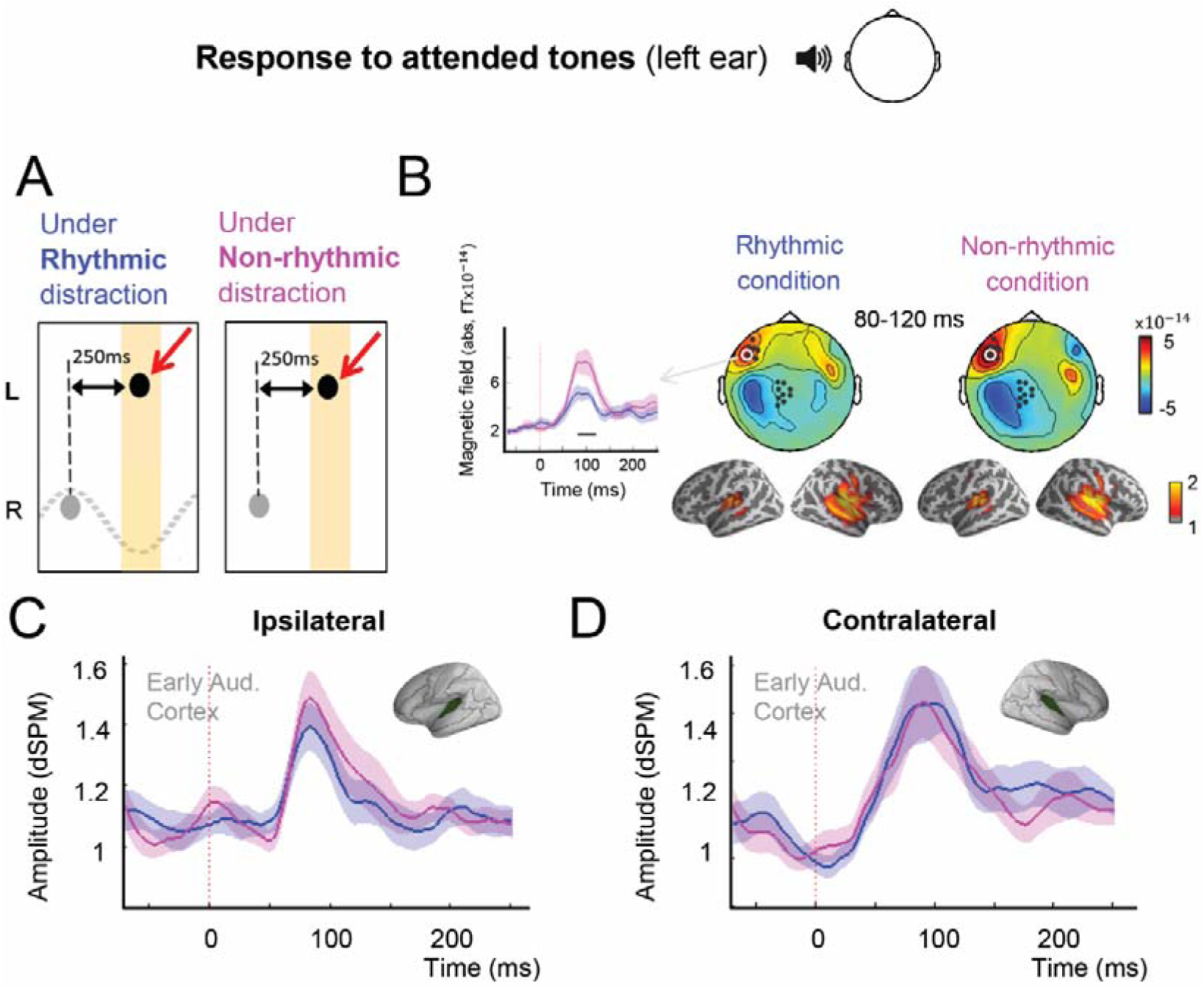
Evoked response locked to attended non-coinciding tones, presented to the left ear. (A) An illustration of the temporal position of these attended tones (black, marked with a red arrow) relative to adjacent distractor tones (gray). (B) Top: Scalp topographies of the M1 response to attended tones under Rhythmic (left) and Non-rhythmic (right) distraction. Black dots indicate MEG sensors where there was a significant difference in the ERF between the Rhythmic and Non-rhythmic distraction conditions (α =0.05, cluster corrected). The time-course of response from one example sensor is shown, comparing response in the Rhythmic (blue) vs. Non-rhythmic (pink) distraction conditions. Bottom: the estimated sources of the MEG recorded response to attended tones presented to the left ear. (C) ERF time course evoked by attended tones in Early Auditory Cortex (see inset), under Rhythmic (blue) and Non-rhythmic (pink) distraction, recorded over the ipsilateral (left) hemisphere. Shaded areas denote SEM. (D) Same for the contralateral (right) hemisphere.

#### Event-Related Analysis: Non-Coinciding Distractor Tones

Besides testing how distractors’ rhythmicity affected the responses to attended tones, we also tested its effect on responses to the distractors themselves. Here too, analysis was limited to distractors that did not coincide temporally with attended sounds (appearing exactly 250 ms after the previous tone; Fig. 4A), as they were matched for local acoustics between conditions. The scalp distribution of ERFs to distractor tones did not conform to the typical auditory response (Fig. 4B). Cluster based permutation tests revealed significant difference between the response to Rhythmic vs. Non-rhythmic distractors. Over the contralateral (left) hemisphere, these differences were evident throughout the entire ERF time-course, from 0-240ms (all clusters p<0.005), and over the ipsilateral (right) hemisphere the differences were significant near the distractor onset (0-40ms, p=0.004) and in a later time window betwwen 80-120ms (p<0.002). Source analysis helped disentangle the neural origins of these responses, and pinpoint the effects of distractor rhythmicity to different components over time. Specifically, two separate effects were found at the source-level: First, there was a significant effect during the pre-stimulus baseline period, starting 40 ms prior to stimulus onset, with enhanced baseline for Rhythmic compared to Non-rhythmic distractor tones. This effect was relatively widespread, and mapped onto temporal and parietal areas, including bilateral Early and Association Auditory cortices, bilateral Insular and Frontal-Opercular cortex, right Posterior-Opercular cortex and left TPJ. (Fig. 4C,D; right cluster p<0.001, left cluster p=0.029). In some regions, this enhanced response in the Rhythmic condition persisted up to ∼80ms post stimulus. In addition, a later response ∼150ms was observed, that was reduced for Rhythmic vs. Non-rhythmic distractors, primarily in Early auditory cortex in the right hemisphere (cluster p=0.005) with a similar trend in Early and Association auditory cortex in the left hemisphere (cluster p=0.067).

**Figure 4:**
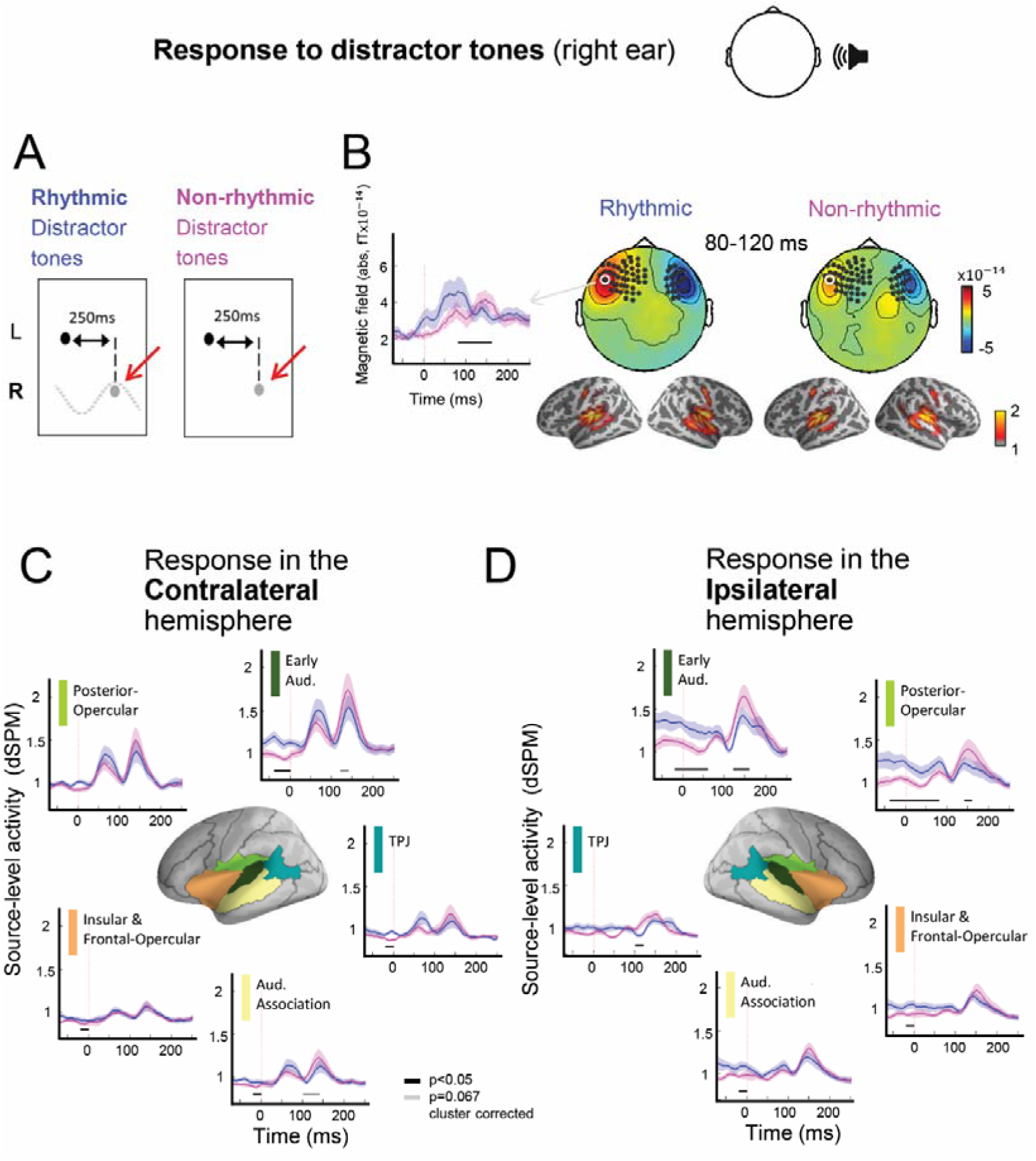
Evoked response locked to distractor non-coinciding tones, presented to the right ear. (A) An illustration of the temporal position of these distractor tones (gray, marked with a red arrow) relative to adjacent attended tones (black). (B) Top: Scalp topographies of the M1 response to attended tones under Rhythmic (left) and Non-rhythmic (right) distraction. Black dots indicate MEG sensors where there was a significant difference in the ERF between the Rhythmic and Non-rhythmic distraction conditions (α=0.05, cluster corrected). The time-course of response from one example sensor is shown, comparing response in the Rhythmic (blue) vs. Non-rhythmic (pink) distraction conditions. Bottom: the estimated sources of the MEG recorded response to distractor tones presented to the right ear. (C) ERF time courses evoked by distractor tones in predefined cortical regions, under Rhythmic (blue) and Non-rhythmic (pink) distraction, recorded over the contralateral (left) hemisphere. Shaded areas denote SEM. Horizontal lines indicate time-windows with significant differences between the conditions (Black: p<0.05, Gray: p=0.067, cluster corrected. (D) Same for the ipsilateral (right) hemisphere.

## Discussion

The current study was aimed at testing whether temporal predictability of distractor sounds is utilized to benefit selective attention, and the underlying neural mechanisms. We focused specifically on rhythmic distractors, given their ecological relevance and the established contribution of rhythms to Dynamic Attending Theory (Nobre et al. 2007; Haegens and Zion Golumbic 2018; Jones 2019). We were curious whether the principles of Dynamic Attending Theory, namely that predicting *when* a stimulus will occur facilitates perception, also extend to stimuli that should be ignored. We find that, indeed, target-detection accuracy improved when it was possible to predict that no distractor would occur, i.e., at the off-phase of a distracting rhythm. Moreover, neural responses to both distractor and attended tones were modulated by the temporal structure of the distractors. For Rhythmic distractors, activity in bilateral auditory cortex and temporo-parietal regions was modulated relative to Non-rhythmic distractors during both the pre-stimulus period as well as in response to the sounds themselves. Neural responses to attended tones were also affected by distractor rhythmicity, as manifest by enhanced SSR in contralateral auditory cortex, accompanied by slightly reduced evoked response in ipsilateral auditory cortex. These combined behavioral and neural findings support the proposal that not only are temporal predictions formed for task-irrelevant sounds, but that these predictions are utilized to reduce their interference and promote selective attention.

### Effects of Distractor Rhythmicity on Performance

One way to assess the effect of distractor rhythmicity is to look at whether performance on the main task was modulated by the type of distractor present. When looking at the average performance collapsed across all target tones, we found no main effect of distractor rhythmicity. However, when separating targets according to their proximity to distractor sounds, we find significant improvement for targets falling on the *off-phase* of a distracting rhythm. These are moments that not only was the target predictable, but additionally it was possible to predict that *no distractor* would occur.

Several previous behavioral studies have also noted improved performance on auditory tasks in the presence of Rhythmic vs. Non-rhythmic distractors (Devergie et al. 2010; Andreou et al. 2011; Rimmele et al. 2012). However, in those studies, the attended stimuli themselves were not necessarily predicable. In the current design, attended stimuli were presented at a fixed rate and hence perfectly predictable, which according to Dynamic Attending Theory should already render them optimal for processing. Nonetheless, the boost in performance for non-coinciding targets suggests that the temporal predictability of distractor sounds carry an added value for perception above and beyond the predictability afforded by the attended stimuli themselves. Additive effects of temporal predictions from multiple sources and at different time-scales have previously been reported for subthreshold audio-visual interactions (ten Oever et al. 2014), however in that study both types of predictive cues were attended. The finding that temporal predictions are formed for task-irrelevant sound and affect performance on the attended task suggests that the principles of Dynamic Attending Theory may extend to ignored sounds. According to this perspective, just as processing resources are dynamically allocated to points in time when attended sounds are expected (Ro et al. 2003; Nobre et al. 2007; Rohenkohl et al. 2012; Lange 2013; ten Oever et al. 2017; Haegens and Zion Golumbic 2018; Auksztulewicz et al. 2019; Jones 2019), they can similarly be dynamically withdrawn from points in time when distractors are expected (Horton et al. 2013; Gray et al. 2015). This perspective gains support from the way neural responses to the distractor sounds themselves were modulated by their temporal structure, which we now turn to discuss.

### Neural Encoding of Rhythmic Distractors

Neural responses to distractor tones were affected by their temporal structure in two ways. The first is elevated activation in the **pre-stimulus period** ∼40 ms prior to the onset of Rhythmic vs. Non-rhythmic distractors. This baseline-shift was observed bilaterally in temporal regions, including early and association auditory cortex, and extended anteriorly towards the insula, as well as posteriorly to the TPJ (in the left contralateral hemisphere only). Given our strict control of local low-level acoustics surrounding the non-coinciding distractors, this baseline shift can reliably be attributed to the difference in global temporal structure of the Rhythmic vs. Non-rhythmic distractors. Modulation of pre-stimulus baseline activity arguably reflect anticipatory effects, brought about when the timing of upcoming stimuli can be predicted. Similar effects are observed when the time of upcoming stimuli is either explicitly cued (Sanders and Astheimer 2008; Chennu et al. 2013; Breska and Deouell 2016, 2017; Mento 2017) or can be inferred from temporal regularities within the stimulus (Sohoglu and Chait 2016; ten Oever et al. 2017; Auksztulewicz et al. 2019). Mechanistically, these anticipatory effects may be attributed to entrainment of intrinsic oscillations so as to increase excitability at specific points in time (Schroeder and Lakatos 2009; Zoefel et al. 2018; Lakatos et al. 2019) or to the buildup of expectation through non-oscillatory mechanisms (e.g. the contingent negative variation ERP response; Pfeuty et al. 2005; Breska and Deouell 2017; Novembre and Iannetti 2018). Distinguishing between these two alternatives is beyond the scope of the current study. However, regardless of the underlying mechanism, the baseline modulation observed here for distractor tones provides strong indication that temporal predictions are formed for rhythmic sounds, even if they are task-irrelevant and presumably outside the ‘focus’ of attention.

Not only were temporal predictions formed in anticipation of rhythmic distractors, but they also affected sensory responses to these sounds, as reflected by modulation of the **evoked response to rhythmic distractors**. ERFs followed a typical biphasic time-course, with an early response peaking at ∼80ms followed by a later response at ∼150ms. The early response was somewhat enhanced for Rhythmic distractors, possibly reflecting a carry-over effect from the baseline shift. However, the pattern reversed for the later response, which was significantly reduced for Rhythmic vs. Non-rhythmic distractors. In contrast to the wide-spread baseline-shift for Rhythmic distractors, this later suppression was more focused to bilateral Early and Association Auditory cortex, and in source space was observed to be more prominent in the ipsi-lateral hemisphere.

One way to interpret the suppression of sensory responses in auditory cortex for rhythmic distractors is within the predictive-coding framework, which posits that evoked response for predictable stimuli are reduced, reflecting a smaller prediction-error (Lange 2009; Bastos et al. 2012; van Atteveldt et al. 2015; Rummell et al. 2016; Sherwell et al. 2017; Friston 2019). According to this interpretation, the effects of predictability on sensory encoding of sounds that are to-be-ignored, are qualitatively similar to previously reported effect of for attended sounds (Sherwell et al. 2017).

However, another interpretation could be that the reduced response to Rhythmic distractors is specifically linked to their status as distractors, and reflects their more effective perceptual suppression due to their temporal predictability. One of the most established operations of selective attention is that evoked responses for task-irrelevant stimuli are suppressed, particularly in dichotic listening paradigms where two stimulus streams are presented concurrently (Hillyard et al. 1973; Woldorff and Hillyard 1991; Näätänen et al. 1992; Ghatan et al. 1998; Kawashima et al. 1999; Ding and Simon 2012; Zion Golumbic et al. 2013). This top-down attentional suppression is thought to involve interactions between sensory cortex and frontal executive control regions (Barbas et al. 2013; Salo et al. 2017; Kuchibhotla and Bathellier 2018), as well as fronto-parietal attentional networks. Several studies have shown that responses to distractors are additionally modulated by how easy/hard they are to ignore (Alain and Woods 1993; Bidet-Caulet et al. 2007; Chait et al. 2010; Noonan et al. 2018), or as listeners become more proficient at ignoring them (Melara et al. 2002b, 2012), although the direction of these effects varies across studies, as does the operationalization of ‘distractor sounds’. In line with the current results, there is evidence from intracranial recordings of cortical responses to two competing sound streams, suggesting that selective attention is facilitated by reducing the representation of irrelevant information in the auditory cortex (Bidet-Caulet et al. 2007). Accordingly, we may interpret the current results as reflecting the utilization of temporal prediction of distractors (formed during the baseline period) in order to actively suppress their sensory encoding more effectively.

These two interpretations are not mutually exclusive, and many also have additive effects. For example, it is possible that responses to Rhythmic distractors are suppressed both because they are predictable AND because they need to be ignored. This theoretical discussion highlights the complex interaction that exists between prediction and attention (Aron 2007; Lange 2013; Heilbron 2018; Press et al. 2019). One way to disentangle their independent contributions would be to compare the effects of rhythmicity on both attended and ignored sounds, an avenue that will be pursued in future research. However, regardless of whether the observed effects are solely due to prediction or also reflect active suppression due to distractor predictability, the current findings clearly demonstrate that the temporal structure of the distractor sounds affects how they are encoded in the brain.

### Effects of Distractor Rhythmicity on Neural Responses to Attended Tones

Not only did distractor rhythmicity affect responses to the distractors themselves, but it affected neural encoding of attended tones as well. Effects of distractor rhythmicity on the evoked response to attended tones were observed when looking at both the SSR and stimulus-locked ERFs. We find increased lateralization of the SSR to the contra-lateral hemisphere and decreased evoked response in the ipsi-lateral hemisphere (although this effect was statistically significant only at the scalp but not at the source level). Both of these effects converge in making the sensory representation of attended sounds more prominent in the contra-lateral auditory cortex.

Auditory responses to unilateral sounds are generally observed in bilateral auditory cortex, due to the early cross over of input in the ascending auditory pathway (Barnes et al. 1943; Alain and Winkler 2012). However, in dichotic-listening conditions, auditory responses become increasingly lateralized (Fujiki et al. 2002; Kaneko et al. 2003; Brancucci et al. 2004; Okamoto et al. 2007; Lazzouni et al. 2010), as also observed here when comparing the SSR to attended tones in the No-distraction condition vs. the two conditions with distractors. In other words, presenting different sounds to each ear de-couples the two auditory cortices, such that input to each ear is primarily encoded in the contra-lateral cortex (Ross et al. 2005). It has been hypothesized that this increased laterality in dichotic conditions may also be useful for selective attention, simplifying focusing spatial attention toward one ear while suppressing input from the task-irrelevant ear. The current results suggest that this spatial biasing was more effective when the distractor sounds were temporally predictable, serving as further indication for the effect of distractor rhythmicity on encoding the attended sounds themselves.

The SSR and ERF are two complementary ways to look at the sensory responses evoked by the attended sounds. We note that although both metrics were modulated by distractor rhythmicity in a manner consistent with contra-lateral bias, the specific manifestations of these effects were not identical. Specifically, when distractors were Rhythmic, the SSR was *enhanced* in the contra-lateral hemisphere but not difference was observed in the ipsi-lateral hemisphere. Conversely, the M1 component of the ERF did not differ between the conditions in the contra-lateral hemisphere, but was *decreased* in the ipsi-lateral hemisphere when distractors were Rhythmic. How should we contend with this difference in response patterns? This ties into a long standing debate about whether the SSR and ERF are two ways of looking at the same sensory response – in the frequency and time domain respectively – or whether they capture different aspects of the sensory response to sounds (Shah et al. 2004; Mazaheri and Jensen 2010; Stupacher et al. 2016; Haegens and Zion Golumbic 2018; Zoefel et al. 2018). Moreover, even if the SSR is merely a frequency-domain representation of rhythmic ERFs, amplitude modulations of individual components within the ERF (e.g. the M1) might not necessarily be reflected in the same way when performing spectral analysis on the entire response, since it is affected by the entire shape of the ERF. It is also worth noting that in the current study, ERFs were analyzed only for a subset of the trials (non-coinciding tones constituted 20-25% of the sounds), which might have also contributed to the differences between the two metrics. However, despite the discrepancy between the specific way that the distractor rhythmicity effect manifested in the SSR and ERFs, the direction of the two effects does converge, emphasizing increased contra-lateral bias for attended sounds when presented on the background of Rhythmic distractors.

### Conclusions

Attention is often thought of as the prioritization of behaviorally relevant inputs over task-irrelevant ones (Desimone and Duncan 1995). However, our results highlight the fact that not all distractors are treated alike. Rather, distractor stimuli themselves may carry information that can be utilized to facilitate selective attention. In particular, when distractors are rhythmic, the system forms predictions about their upcoming timing. These predictions influence sensory responses to the distractors themselves and, importantly, also affect how attended sounds are processed and the behavioral responses to them. Therefore, we suggest that the notion of Dynamic Attending is not limited to utilizing temporal information from task-relevant stimuli, but that the system also makes use of predictabilities in background and task-irrelevant sounds to further sharpen the precision of attention allocation. This work contributes to our growing understanding of the complex relationship between prediction and attention, that serve as key mechanism for active sensing (Aron 2007; Schroeder et al. 2010; Zion Golumbic et al. 2012b; Lange 2013; Heilbron 2018; Press et al. 2019).

## Acknowledgment

Research was supported by the ISF I-Core Center for Excellence 51/11 to EZG, and by BSF grant #2015385 to EZG. We would like to thank Noa Guttman Hacohen for assistance in experiment setup and data collection. Corresponding author: Zion-Golumbic Elana, The Leslie and Susan Gonda (Goldschmied) Multidisciplinary Brain Research Center, Bar-Ilan University, Building number 901, room 412, Ramat-Gan 5290002, Israel, +972 3 7384430.

